# Astrovirus Replication Is Inhibited by Nitazoxanide *In Vitro* and *In Vivo*

**DOI:** 10.1101/797316

**Authors:** Virginia Hargest, Bridgett Sharp, Brandi Livingston, Valerie Cortez, Stacey Schultz-Cherry

**Author notes:** Address correspondence to Stacey Schultz-Cherry,.

## Abstract

Astroviruses (AstV) are a leading cause of diarrhea especially in the very young, the elderly, and immunocompromised populations. Despite their significant impact on public health, no drug therapies for astrovirus have been identified. In this study we fill this gap in knowledge and demonstrate that the FDA-approved broad-spectrum anti-infective drug nitazoxanide (NTZ) blocks astrovirus replication *in vitro* with a 50% effective concentration (EC_50_) of approximately 1.47μM. It can be administered up to 8 hours post-infection and is effective against multiple human astrovirus serotypes including clinical isolates. Most importantly, NTZ reduces viral shed and clinical disease (diarrhea) *in vivo*, exhibiting its potential as a future clinical therapeutic.

**Importance:** Human astroviruses (HAstV) are thought to cause between 2 and 9% of acute, non-bacterial diarrhea cases in children worldwide. HAstV infection can be especially problematic in immunocompromised people and infants where the virus has been associated with necrotizing enterocolitis, severe and persistent diarrhea, as well as systemic and often fatal disease. Yet no antivirals have been identified to treat astrovirus infection. Our study provides the first evidence that nitazoxanide may be an effective therapeutic strategy against astrovirus disease.

## Introduction

Diarrheal disease is the second leading cause of death in children under 5 years of age, with nearly 1.7 billion cases and 525,000 deaths each year (1). Since their discovery in 1975, human astroviruses have consistently ranked among the leading causes of diarrhea worldwide (2). However, human astrovirus infections can range anywhere from asymptomatic to mild diarrhea and even fatal systemic disease (3, 4). Infections in immunocompetent individuals typically present as watery diarrhea that resolves within 1 to 3 days post-infection without the need for hospitalization (2). Astrovirus outbreaks of frequently occur in assisted living facilities, hospitals and child care centers, where the young, elderly, and immunocompromised populations are at risk of persistent diarrhea leading to wasting (5), and extra-gastrointestinal disease that requires medial intervention, including respiratory disease (6–10), and fatal encephalitis, and meningitis (11).

Despite its high prevalence and the risk of severe disease, no vaccines or drug treatments exist for astrovirus. Only oral or parenteral fluids and electrolytes available to prevent and treat dehydration caused by astrovirus-induced diarrhea. In these studies, we provide the first evidence that nitazoxanide (NTZ), an FDA-approved broad-spectrum anti-parasitic and antiviral drug, inhibits the replication of multiple strains of human astrovirus *in vitro* even when administered up to 8 hours post-infection, and reduces viral shed and diarrhea *in vivo*. This work highlights the potential use of NTZ as an effective therapeutic strategy against astrovirus infection.

## Results

### Nitazoxanide blocks astrovirus replication *in vitro* by inhibiting dsRNA production

To identify an effective antiviral drug against astrovirus, human colon carcinoma (Caco-2) cells were infected with a laboratory strain of human astrovirus strain-1(HAstV-1) at a multiplicity of infection (MOI) of 1 before increasing concentrations of foscarnet, ribavirin, acyclovir, nitazoxanide or DMSO (vehicle control) were added 1 hour post-infection (hpi). Viral capsid protein levels were quantitiated at 24 hpi by immunofluorescent microscopy as described (12). Foscarnet, ribavirin, and acyclovir failed to inhibit HAstV-1 replication even at concentrations of 250 μM (Figure 1a). In contrast, NTZ inhibited HAstV-1 replication in a dose-dependent manner with the 5 μM treatment completely blocking virus replication (Figure1a-b). The 50% effective concentration was calculated as approximately 1.47 μM (Figure 1c). Concentrations of NTZ above 5 μM were associated with decreased cell viability as compared to vehicle alone (DMSO) (Figure 1d). Thus, subsequent studies were performed with NTZ at a concentration of 2.5 μM.

**Figure 1.**
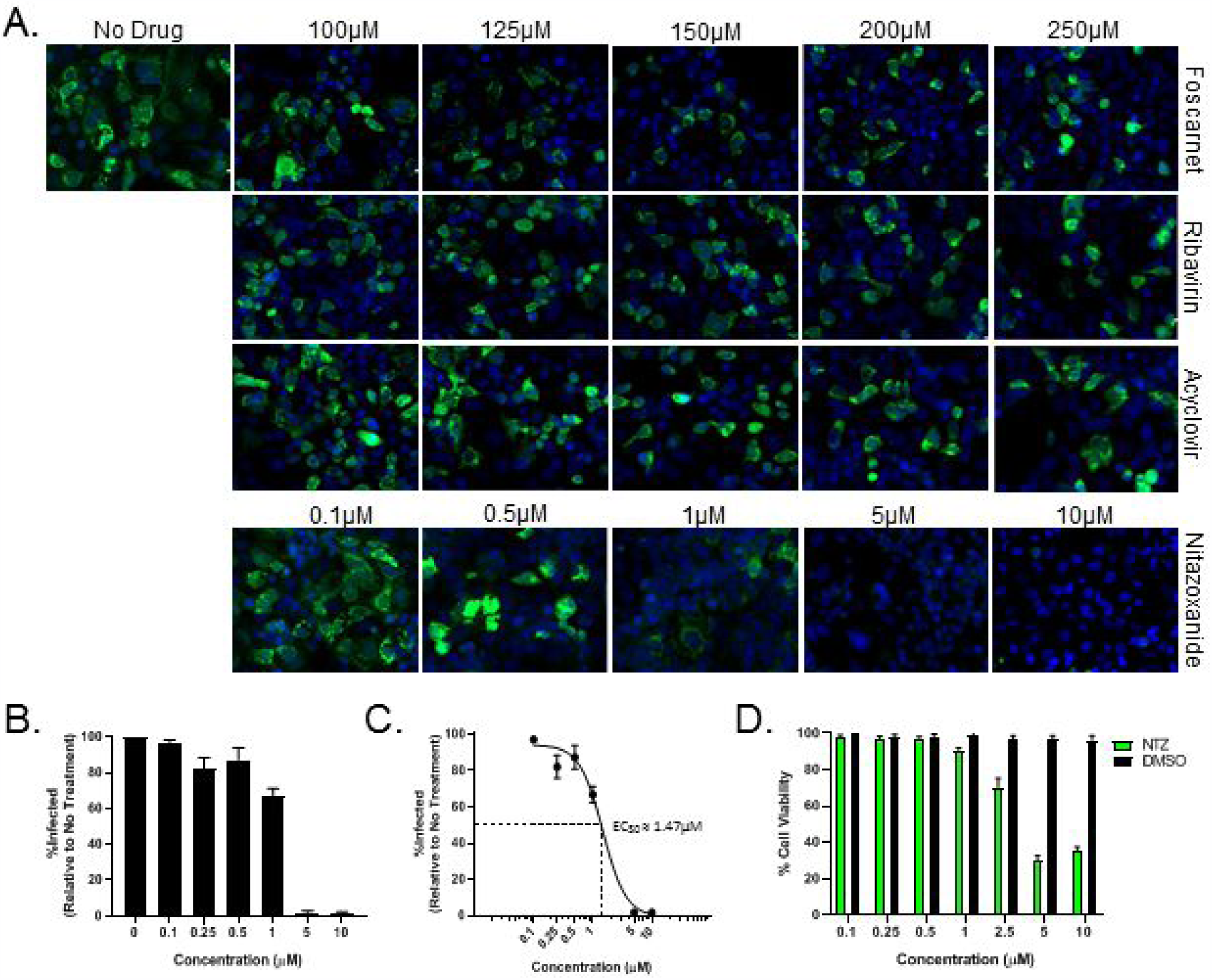
Nitazoxanide inhibits HAstV-1 replication in Caco-2 cells. (A) Caco-2 cells were infected with HAstV-1 at an MOI of 1 and treated with a panel of antivirals (foscarnet, ribavirin, acyclovir, and nitazoxanide) at the indicated concentrations. At 24hpi, cells were fixed and stained with DAPI (blue) and for the presence of astrovirus capsid protein (green). (B) The percent of infected cells was calculated and compared to non-treated cells. (C) Non-linear regression analysis of percent infection data was used to determine the 50% effective concentration (EC_50_). (D) Cell viability of Caco-2 cells following 24 hour treatment with NTZ (green bars) or vehicle alone (DMSO; black bars) at the indicated concentrations was determined by MTT assay. All error bars indicate standard error of the means.

To determine the stage of the viral replication cycle blocked by NTZ, drug was added at 2, 4, 6, 8, and 12 hours after HAstV-1 infection and viral capsid expression quantitated (Figure 2a). Addition of NTZ up to 8hpi completely inhibited HAstV-1 replication (Figure 2b) suggesting it blocks an early stage of the viral life cycle. Indeed, NTZ reduced the formation of double-strand RNA that occurs when the astrovirus genome is generated via its RNA-dependent RNA polymerase (Figure 2c). This method serves as a proxy for early replication given our lack of antibodies to the HAstV-1 non-structural proteins. Since NTZ appeared to exert antiviral activity early in the astrovirus replication cycle, we asked if NTZ activated any innate antiviral pathways. Previous research has shown thiazolides, the active form of NTZ, up-regulate type I and II IFN (13), HAstV has already been shown to be sensitive to IFN treatment, where exogenous addition of IFNβ limited astrovirus replication in a dose-dependent manner (14). We treated Caco-2 cells with NTZ and looked for up-regulation of type I and III interferons (IFN). At 4 hours post NTZ treatment, the shortest amount of time in which NTZ still inhibits HAstV replication, IFNα, IFNβ, or IFNλ were not significantly up-regulated compared to non-treated cells (Figure 2f). This suggests that the induction of IFN is not the mechanism by which NTZ blocks HAstV replication.

**Figure 2.**
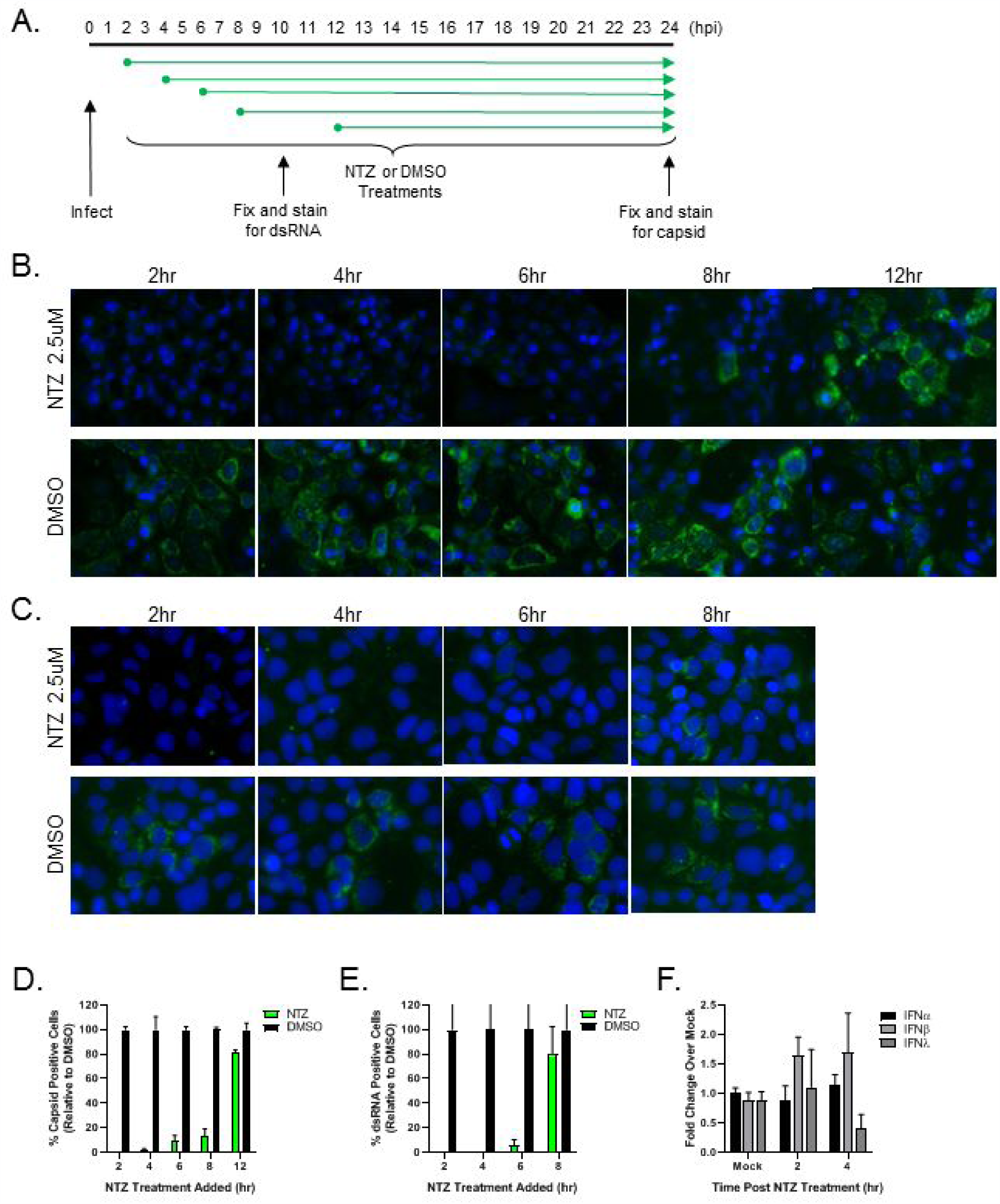
Nitazoxanide inhibits HAstV-1 replication in vitro when added up to 8 hpi. Caco-2 cells were infected with HAstV-1 and at the various times post-infection 2.5μM NTZ or vehicle alone (DMSO) was added as indicated by the schematic in panel A. (B) At 24hpi, cells were fixed and stained with DAPI (blue) and for the presence of astrovirus capsid protein (green). (C) At 10hpi, cells were fixed and stained with DAPI (blue) and for the presence of dsRNA (green). (D) Quantification of the percent of cells with capsid staining from panel B. (E) Quantification of the percent of cells with dsRNA staining from panel C. (F) Real-time RT-PCR for IFNα, IFNβ, and IFNλ was performed on RNA was collected from Caco-2 cells treated with 2.5μM NTZ and normalized to GAPDH. Results are shown as fold increase over untreated cells and error bars indicate standard error of the means.

Finally, we asked if NTZ was effective against multiple HAstV serotypes and clinical isolates. Briefly, Caco-2 cells were infected with four different lab-adapted HAstV serotypes (HAstV-1, 2, 6 and 8) and four clinical isolates SJ177.110 (HAstV-2), SJ60.212 (HAstV-8), SJ88123.E120 (HAstV-1), and SJ88027.E259 (HAstV-1) obtained from patient samples and treated with 2.5 μM NTZ or DMSO (vehicle control). NTZ completely inhibited all HAstV isolates suggesting it is broadly protective against multiple HAstV serotypes (Figure 3a). Within the past decade, novel HAstV more closely related to animal AstV and associated with severe extra-gastrointestinal symptoms have been discovered. To determine if NTZ could also inhibit these non-classical HAstV, we infected Caco-2 cells with VA1, treated the cells with NTZ, and at 24hpi stained for dsRNA. Again, NTZ completely inhibited VA1 replication (Figure 3b). Thus, NTZ exhibits compelling potential as a treatment option across HAstV serotypes and possibly individuals experiencing extra-gastrointestinal symptoms of astrovirus infection.

**Figure 3.**
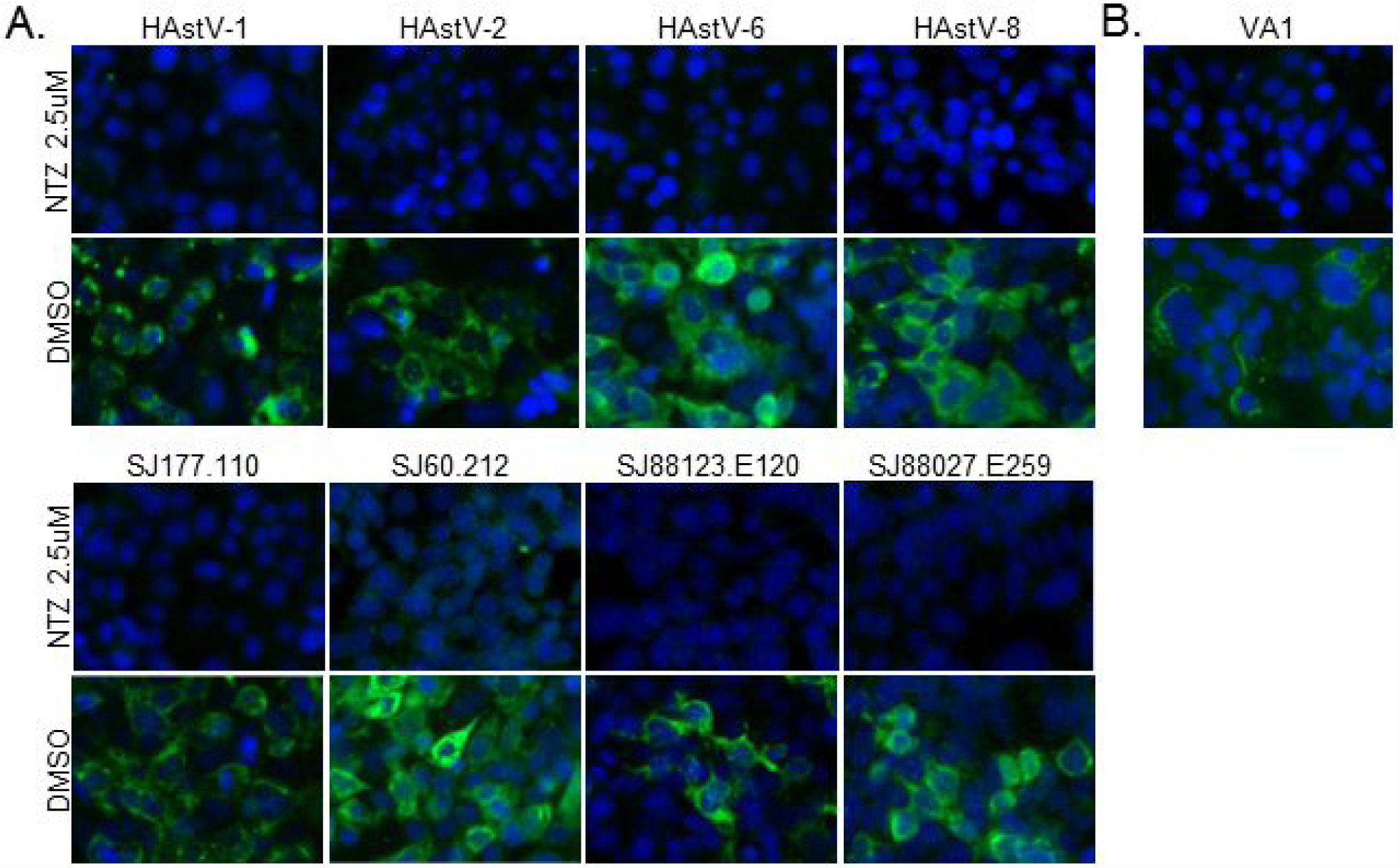
Nitazoxanide inhibits the replication of multiple serotypes and clinical isolates of human astrovirus. (A) Caco-2 cells were infected with lab adapted virus serotypes (upper panels) or clinical isolates (lower panels) and treated with 2.5μM NTZ or vehicle alone (DMSO). At 24hpi, cells were fixed and stained with DAPI (blue) and for the presence of astrovirus capsid protein (green). (B) Caco-2 cells infected with VA1 were fixed at 24hpi and stained with DAPI (blue) and for the presence of dsRNA (green).

### NTZ reduces viral replication and clinical disease in *vivo*

To determine if NTZ reduced disease, we used the only small animal model exhibiting astrovirus-induced diarrheal disease, turkey poults (15). Briefly, 5 day old turkey poults were orally gavaged with 100 mg/kg NTZ once daily for 4 days prior to oral infection with intestinal filtrate containing between 10^12^-10^13^genome copy units turkey astrovirus (TAstV-2) in 500 μl PBS and 3 days post-infection (Fig 4a). Stool samples were scored daily by four blinded volunteers as previously described (16). The scoring scale ranged from 1 to 4 based on color and consistency, with scores of 3 and 4 being considered diarrhea. Stool was also collected to quantitate viral titers. Fewer NTZ-treated poults had clinical disease (Fig 4b) and these poults had significantly less virus in their stool throughout the course of infection (Fig 4c). The poults showed no adverse symptoms from receiving NTZ alone. Excitingly, this gives evidence NTZ may be an effective therapeutic for astrovirus-induced diarrheal disease in patients.

**Figure 4.**
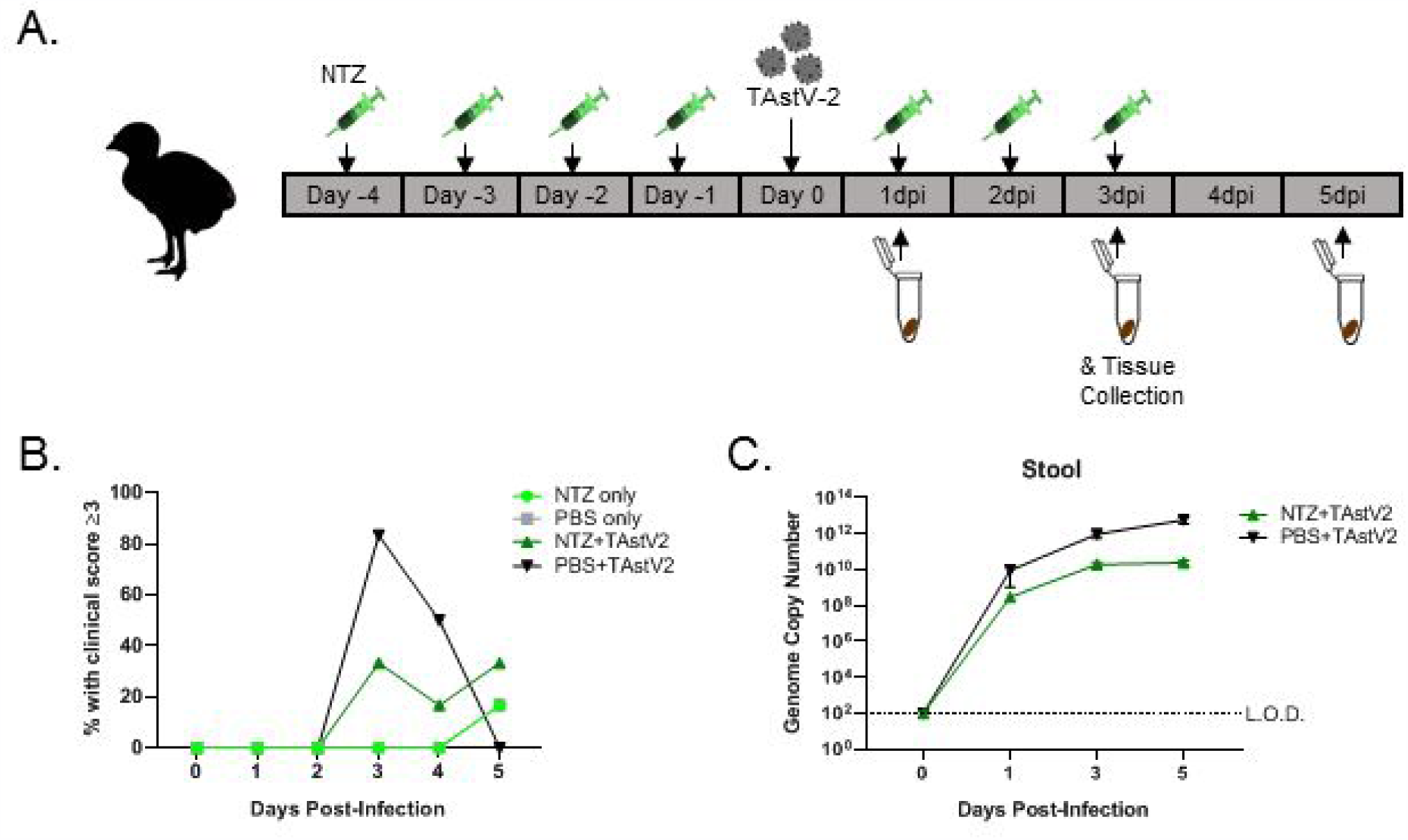
Nitazoxanide reduces clinical symptoms and viral titers in turkey poults. (A) Turkey poults (n=6/group) were infected with turkey astrovirus (TAstV-2) from intestinal filtrate. Four days prior to infection and three days post-infection poults were treated with NTZ. Poults were monitored for clinical score daily and stool was collected to measure viral RNA titer every other day. (B) Percentage of poults with clinical scores of 3 or higher in the groups: NTZ treatment alone (green circle), no antiviral treatment (gray square), TAstV-2 infected without antiviral treatment (black triangle), and TAstV-2 infected with NTZ treatment (green triangle). (C) Viral RNA titer of stool collected from infected poults with NTZ treatment (green triangles) or no antiviral treatment (black triangles). All error bars indicate standard error of the means, and dashed line represents the limit of detection.

## Discussion

Our study provides the first evidence of an effective antiviral for astrovirus infection. We showed NTZ is broadly protective against multiple HAstV serotypes and reduces the production of dsRNA during infection *in vitro* with an EC_50_ of approximately 1.47 μM. Additionally, we showed the potential NTZ has as a clinical therapeutic by its ability to reduce viral shed and clinical disease in our symptomatic turkey poult model.

Nitazoxanide, 2-acetyloxy-N-(5-nitro-2-thiazolyl) benzamide (Alinia; Romark Laboratories), is a thiazolide compound for treatment of both intestinal protozoal and helminthic infections specifically *Giardia lamblia* and *Cryptosporidium parvum* (17). Recently this compound has been shown to have antiviral properties as well. The use of NTZ in vitro has been reported as an antiviral against influenza virus (18), rotavirus (19), norovirus (20), Japanese encephalitis virus (JEV) (21), rubella virus (22), Zika virus (23), hepatitis C virus (24), and hepatitis B virus (25). Successful clinical trials have demonstrated its effectiveness in treating influenza (26), norovirus and rotavirus (20, 27, 28), hepatitis B virus (29), and hepatitis C virus (24, 30). Its mechanism of action against protozoa is due to its interference with pyruvate:ferredoxin oxidoreductase (PFOR) enzyme-dependent electron transfer reactions (31). While its antiviral action is currently unknown, research suggests it may be through the induction of the interferon response via activation of protein kinase R, or disruption of the unfolded protein response (17). We show that NTZ disrupts astrovirus infection early in the replications cycle causing a significant decrease in the production of dsRNA. The inhibition by NTZ at an early stage of infection was also seen with JEV (21). Recent research has shown thiazolides upregulate type I and II IFN (13), which modulate the immune system and could be how NTZ creates a broadly antiviral state. However, the rapid kinetics with which NTZ inhibits HAstV replication (Figure 2) suggests that the induction of IFN is not responsible.

Astroviruses are classified into genotypes, but within the classical human genotype, *Mamastrovirus 1 (MAstV1)*, strains are further divided into serotypes (HAstV-1-8) based on their antigenicity and genetic differences in the complete capsid sequence (2). These genetic differences between astrovirus serotypes can confer differences in replication kinetics and symptom severity (32). Thus, finding a compound that broadly inhibits astroviruses across genotypes and serotypes is crucial. We found that NTZ is broadly protective across multiple HAstV serotypes, including the dominant strain worldwide, HAstV-1 (33), and patient isolates. Excitingly, NTZ shows efficacy against at least one non-classical HAstV genotype, *MAstV9*, specifically the VA1 serotype (Figure 3b). Non-classical HAstV (*MAstV6, 8, and 9*) have been linked to severe extra-gastrointestinal symptoms. To date, the non-classical HAstV genotypes have been associated with eight cases of encephalitis or meningitis (11), with VA1 being identified in five of those cases. Since the 1980’s, the incidence of classic HAstV has been declining (5), studies have shown seroprevalence of VA1 and MLB1 is 65% (34) and 86% (35), respectively, and could account for the displacement of circulating classic HAstV. Testing the susceptibility of primary astrovirus isolates as well as non-classical HAstV to NTZ increases our confidence that this drug would be effective in a clinical setting against circulating strains of HAstV.

Turkey poults exhibit age-dependent diarrhea similar to humans when infected with TAstV making them the only clinically relevant small animal model for astrovirus identified to date (15, 36). We found that NTZ reduced virus levels shed in stool. We found a significant decrease in stool viral titers with NTZ-treated poults having nearly 2 logs less virus at 5 days post-infection. We also showed that viral titers began to plateau in the NTZ-treated poults at 5 days post-infection while untreated poults still showed increasing titers. This suggests NTZ treatment may lead to faster clearance of the virus, however additional studies taken out further are needed to definitively prove this.

These studies were repeated, however, due to the seasonality of turkey breeding and limited availability of poults, a different breed of turkey, royal palm, was used. With the royal palm poults, we again saw a reduction of viral titer in the stool. Additionally, we tested the small intestinal tissue to quantitate viral RNA. The reduction in stool titers was recapitulated in the tissue, where NTZ-treated poults had 2 logs lower virus in the duodenum, and about 1 log lower virus in both the jejunum and the ileum (data not shown). However, NTZ-treatment had no effect on the reduction of clinical symptoms in the royal palm poults. The mechanism by which TAstV-2 induces diarrhea could be why this reduction was not statistically significant throughout infection. We know that administration of capsid alone is sufficient to induce diarrhea in turkey poults (16). From our *in vitro* work we believe NTZ blocks replication around the point where the AstV genome is copied. Therefore, NTZ treatment may not be able to fully inhibit AstV-induced diarrhea. In addition to this point, we administered between 10^12^-10^13^ genome copy units to each poult. While there have been reports of virus shed in humans at this level (32), it is a large viral dose that may not be representative of natural infection. This work provides the first evidence that NTZ may be an effective antiviral option against a broad range of HAstV, including both classical and non-classical genotypes and limits viral titers *in vivo*.

## Materials and Methods

### Cells and Virus Propagation

The human intestinal adenocarcinoma cell line Caco-2 was obtained from ATCC (HTB-37). Cells were propagated in minimum essential medium (MEM; Corning) supplemented with 20% fetal bovine serum (FBS; Benchmark), GlutaMax-I (Gibco), 1 mM sodium pyruvate (Gibco), and penicillin-streptomycin (Gibco).

Lab adapted human astrovirus stocks (HAstV-1, HAstV-2, HAstV-6, HAstV-8) were propagated in Caco-2 cells, and the titer of the viruses were determined on Caco-2 cells by the fluorescent-focus assay (focus-forming units [FFU]) as previously described (12).

Clinical isolates (SJ177.110, SJ60.212, SJ88123.E120, and SJ88027.E259) were derived from remnant fecal samples submitted for clinical diagnostic testing at St. Jude Children’s Research Hospital. All samples were de-identified before testing. The St. Jude Institutional Review Board approved this study with a waiver of consent. All isolates were propagated in Caco-2 cells. Briefly, a 10-20% dilution of stool extract, positive for HAstV by RT-PCR, was filtered through a 0.22μm filter. The extract was diluted 1:10 in MEM + 5μg/ml porcine trypsin before adsorption onto Caco-2 cell monolayers. Following a 1 hour adsorption period at 37°C, the inoculum was removed and replaced with MEM containing 10μg/ml porcine trypsin and 0.3% BSA. The titer of the viruses were again determined on Caco-2 cells by the fluorescent-focus assay (12). TAstV-2 stocks were prepared from intestines collected from infected turkey poults. Briefly, pieces of intestine were suspended in 0.5 ml PBS in multiple tubes, homogenized using 2-mm zirconium oxide beads (Next Advance) beads for 4 minutes on speed setting 4 (Next Advance air cooling bullet blender), and pelleted by centrifugation at 12,000 rpm for 5 minutes. The supernatants were pooled and filtered through a 0.2-μm filter (fecal filtrate), and viral copy number was quantified by real-time RT-PCR.

### *In vitro* HAstV Infection

Briefly, 5 × 10^4^ cells were seeded into 96-well tissue culture plates (Corning), and after 2 days, the cells were inoculated with virus (HAstV-1, clinical isolates, VA1) in serum-free MEM for 1 hour at 37°C, at which time the virus was replaced with MEM containing 0.3% BSA and infection was allowed to proceed until 24hpi unless otherwise stated.

NTZ treatment was carried out in serum-free MEM and added following the 1 hour virus adsorption period unless otherwise stated in the experimental design.

### Immunofluorescent Staining

Cells were fixed with 100% ice-cold methanol for 15 minutes, and then blocked with 5% normal goat serum (NGS; Gibco) in PBS at room temperature. The cells were stained with HAstV mouse monoclonal antibody 8E7 (2 μg/ml DakoCytomation) for 1 hour at room temperature followed by anti-mouse IgG labeled with Alexa Fluor 488 (anti-mouse IgG-Alexa Fluor 488; Invitrogen) secondary antibodies and 4′,6′-diamidino-2-phenylindole (DAPI; Sigma) for 30 minutes at room temperature. Staining was imaged on EVOS® FL Cell Imaging System and analyzed using ImageJ 1.50i software.

### MTT Cell Viability Assay

Cell viability was tested using an MTT Cell Proliferation assay kit (Abcam) according to the manufacturer protocol. Briefly, cells were treated with varying concentrations of nitazoxanide in serum free media for 24 hours. The nitazoxanide containing media was removed and replaced with a 50:50 mixture of MTT reagent and serum free media. The cells were incubated with the mixture at 37°C for 3 hours. Following incubation, an MTT solvent solution was added and the plate was placed on an orbital shaker for 15 minutes. The absorbance was then measured at OD595. Cell viability was calculated as a percentage of non-treated cells.

### Animals and NTZ treatment

Broad-breasted white turkey poults were obtained from a commercial hatchery. Five-day-old poults were randomly assigned to groups (n = 6 per group) and housed in individual, temperature-controlled Horsfall units with HEPA-filtered inlet and exhaust air valves, where they were given free access to water and routine turkey starter feed. Poults were orally inoculated with 500μl of TAstV-2 intestinal filtrate, containing approximately 10^12^-10^13^ genome copies, or PBS alone. Stool from individual birds was scored from 1 to 4. Scoring was performed daily post-infection. Scores of 3 (liquid or loose stool with some undigested food or solid material) and 4 (watery stool with no solids present) were defined as diarrhea, in accordance with previously published work from Meliopoulos et al. (16).

For NTZ treatment, poults were orally administered 100mg/kg nitazoxanide in 500ul of ultrapure water. Administration of NTZ was carried out 4 days prior to infection and 3 days postinfection.

### Turkey Astrovirus qRT-PCR assay

TAstV-2 genome copies were determined as previously described (16). Briefly, viral RNA was isolated from 10% stool by the MagMAX-96 AI/ND Viral RNA Isolation Kit (Applied Biosystems) according to the manufacturer’s protocol. PCR was performed on 3 µl of each sample using TaqMan™ Fast Virus 1-Step Master Mix (Applied Biosciences) with 600nM forward primer 5′GACTGAAATAAGGTCTGCACAGGT, 600nM reverse primer 5′AACCTGCGAACCCTGCG, and 200nM probe 6-carboxyfluorescein (6FAM)-ATGGACCCCCTTTTTCGGCGG-BHQ1 (black hole quencher) under the following conditions: 50°C for 5 min, 95°C for 20 s, followed by 45 cycles, with one cycle consisting of 95°C for 3 s and 60°C for 30 s on a Bio-Rad CFX96 real-time PCR detection system. The number of genome copies/µL of total RNA was determined using a standard curve generated from a synthesized TAstV-2 DNA from nucleotides 4001 to 4201 with a known copy number (calculated using Thermo Fisher Scientific DNA Copy Number and Dilution Calculator). Log_10_ dilutions of the synthesized TAstV-2 DNA were used for real-time RT-PCR as described above.

## Acknowledgments

We thank Rebekah Honce, Pamela Freiden and Dr. Victoria Meliopoulos for their expert poop scoring abilities; Sean Offord and Sharon Lokey from the St. Jude Animal Resources Center for assistance with turkey studies; and Dr. David Wang for graciously providing VA1 for these studies.

These studies were funded by National Institute of Allergy and Infectious Diseases R21 AI135254-01, and ALSAC.

